# A universal approach for integrating super large-scale single-cell transcriptomes by exploring gene rankings

**DOI:** 10.1101/2021.08.23.457305

**Authors:** Hongru Shen, Xilin Shen, Mengyao Feng, Dan Wu, Chao Zhang, Yichen Yang, Meng Yang, Jiani Hu, Jilei Liu, Wei Wang, Yang Li, Qiang Zhang, Jilong Yang, Kexin Chen, Xiangchun Li

## Abstract

Advancement in single-cell RNA sequencing leads to exponential accumulation of single-cell expression data. However, there is still lack of tools that could integrate these unlimited accumulation of single-cell expression data. Here, we presented a universal approach *iSEEEK* for integrating super large-scale single-cell expression via exploring expression rankings of top-expressing genes. We developed *iSEEEK* with 13.7 million single-cells. We demonstrated the efficiency of *iSEEEK* with canonical single-cell downstream tasks on five heterogenous datasets encompassing human and mouse samples. *iSEEEK* achieved good clustering performance benchmarked against well-annotated cell labels. In addition, *iSEEEK* could transfer its knowledge learned from large-scale expression data on new dataset that was not involved in its development. *iSEEEK* enables identification of gene-gene interaction networks that are characteristic of specific cell types. Our study presents a simple and yet effective method to integrate super large-scale single-cell transcriptomes and would facilitate translational single-cell research from bench to bedside.

## Introduction

Large volume of single-cell transcriptomes is accumulating rapidly. Technical improvements in single-cell RNA sequencing (scRNA-seq)^1^ lead to rapid drop in sequencing cost and allows for millions of cells to be sequenced. This was exemplified by the establishment of international collaborative projects on single-cell such as Human Cell Atlas^2^, COVID-19 Atlas^3^, Single Cell Expression Atlas^4^, Tabula Muris Atlas^5^ and Mouse Cell Atlas^6^, which aim at depicting reference map of single-cell signatures. Consequently, integration of these super large-scale data is a challenge and crucial in the era of single-cell data science^7^.

Traditional single-cell transcriptome analysis methods such as Seurat^8,9^ and Scanpy^10^ are to learn feature representation of gene expression profiles via dimensional reduction on expression profiles of high variable genes (HVGs). While the deep learning methods such as scVI^11^ and MARS^12^, in essence analogous to traditional methods, are to perform dimensionality reduction on gene expression of single-cells specifically in a nonlinear manner. However, there remain several challenges for single-cell analysis. For instance, there are high discrepancies in the selection of HVGs among different methods^13^ and the batch effect further complicates HVG selection^14^. Noise and batch effect are unavoidable as sequencing samples were often compiled from multiple experiments, handling by different personnel, sequenced with different instruments and protocols^15,16^. The batch effect masks the biological variations and entails batch correction. However, overcorrection is often inevitable ^17^.

Herein, we introduced *iSEEEK*, a universal approach for integrating super large-scale single-cell transcriptomes via exploring the rankings of top-expressing genes. We hypothesize that the expression information of a single-cell is manifested by the rankings of its top-expressing genes. Therefore, we formulated feature representation of single-cell transcriptomes as natural language processing (NLP) task in that the sentence of each single-cell was constructed by concatenation of gene symbols of top-expressing genes ordered by their expression levels. Tremendous progress and enormouse achievement were obtained in NLP task. The emergence of GPT^18^, BERT^19^, and ERINE^20^ algorithms revolutionized deep learning in domain of natural language understanding such as document classification, question answering and semantic similarity assessment *etc*. The essence of these algorithms is devoted to modeling associations among tokens and sentences as pretraining task. We developed *iSEEEK* to model the rankings of top-expressing genes on a dataset of 13.7 million single-cells. Subsequently, we applied the pretrained *iSEEEK* in downstream tasks such as delineation of cell clusters on three heterogeneous datasets such as peripheral blood mononuclear cells^9^, Human Cell Atlas^21^ and expression profiles of 20 organs from Tabula Mursi^5^. We also tested the transferability of *iSEEEK* on a new dataset that was not involved in its development. In addition, we demonstrated the applicability of *iSEEEK* to extract gene-gene interaction networks that are specific for CD4/8+ T cells obtained from fluorescence-activated cell sorting (FACS). *iSEEEK* would facilitate the integration of super large-scale single-cell transcriptomes and translational single-cell research from bench to bedside.

## Results

### *iSEEEK*:<u>i</u>ntegration of <u>S</u>ingle-cell <u>E</u>xpression via <u>E</u>xploring <u>E</u>xpression ran<u>K</u>ings of top-expressing genes

*iSEEEK* was trained with masked language model task to model the expression rankings of the top-expressing genes. *iSEEEK* was trained with 13,702,899 single-cells collected from public databases covering a variety of cell types from different human tissues under different conditions and mouse tissues (**Supplementary Table 1**). *iSEEEK* takes as input a sequence of gene symbols ranked by their expression levels (See **Methods**). The model learns the information of the ranking of the *n* top-expressing genes in a decreased order per cell. In this study, we examined *iSEEEK* with the rankings of the top 126 expressing genes. *iSEEEK* was trained as a masked language modeling task^19,22^. In this study, the masked language model task randomly masks some of genes in the input and predict the vocabulary indexes of masked genes based on their bidirectional contexts. The vocabulary consists of 20,706 protein-encoding genes. *iSEEEK* benefits from multi-head self-attention mechanism and bidirectional encoder representation. The aggregation of feature representations from multi-head attentions improved efficiency and precision. We applied the same data sampling strategy during training as proposed by Devlin J. and colleagues^19^: the training data generator randomly chooses 15% of the gene positions for prediction. If the i^th^ gene is chosen, we replace the it with (1) the [MASK] token 80% of the time, (2) a random gene 10% of the time, (III) the original unchanged gene 10% of the time. *iSEEEK* was trained by cross-entropy loss by comparing its predictions to the original genes (**Figure 1A**). *iSEEEK* consists of 8 transformer layers each with 576 hidden units and 8 attention heads. Detailed parameters of *iSEEEK* were listed in **Supplementary Table 2**. The developed *iSEEEK* is able to learn the representations of expression-based gene rankings. The latent features extracted from the pretrained *iSEEEK* model can be used as input for downstream task including delineation of cell clusters, identification of marker genes and exploration of cell developmental trajectory *etc* (**Figure 1B**).

**Figure 1.**
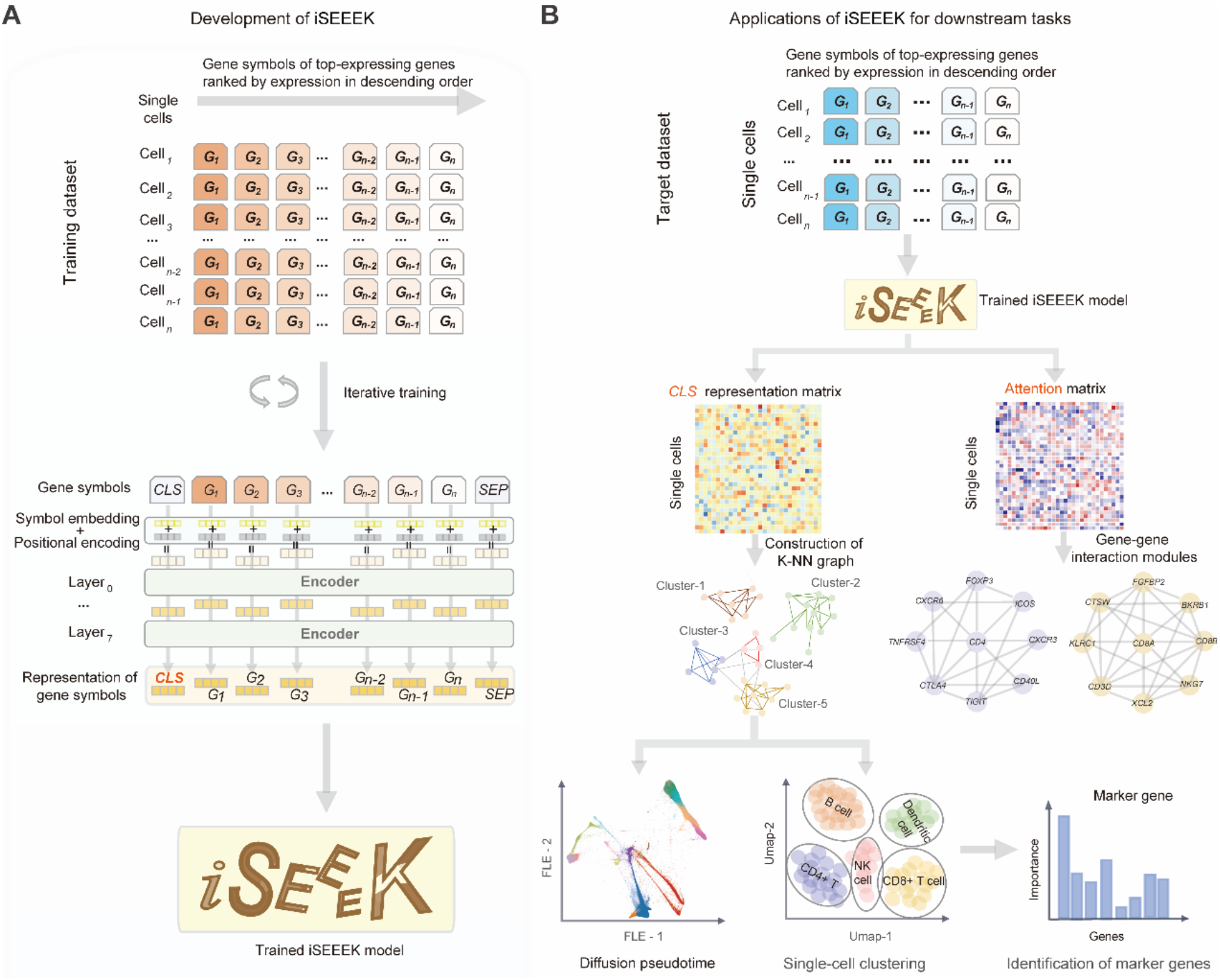
A flowchart depicting the development and downstream applications of *iSEEEK*. (**A**) Development of *iSEEEK* based on the genes symbols of top-expressing genes ranked by expression in descending order for large-scale single-cells. (**B**) Downstream application of *iSEEEK* includes delineation of single-cell clustering, pseudotime inference of cell trajectory, identification of marker genes and exploration of cluster-specific gene-gene interaction modules.

### Clustering performance of *iSEEEK*

We evaluated the clustering performance of *iSEEEK* on three heterogeneous datasets that encompassed bone marrow dataset from Human Cell Atlas Census of Immune Cells^21^ (HCA, n=282,558), peripheral blood mononuclear cells^9^ (PBMC, n=43,073) and Tabula Mursi dataset^5^ (n=54,865 cells). The HCA bone marrow dataset consisted of 18 cell types with different proportions. The PBMC dataset consisted of CD4+ T cell, CD8+ T cell, NK cells, FCGR3A+ and CD14+ monocytes. The Tabula Mursi dataset included single-cells of 20 organs from *Mus musculus*.

*iSEEEK* was able to reveal distinct cell clusters underlying the composition of each dataset. On the HCA bone marrow dataset, the cell subsets were well separated and the megakaryocytes with low proportion (0.32%) were captured by *iSEEEK* (**Figure 2A**). On the PBMC dataset, *iSEEEK* revealed 23 cell clusters involving eight immune cell subgroups (**Figure 2B**). The cytotoxic lymphocyte cells were gathered together but divided into CD4+ T cell, CD8+ T cell and NK cell subgroup, and monocytes with different markers (FCGR3A+ or CD14+) are also well mapped in particular. On the Tabula Mursi dataset from *Mus musculus* composed of 20 mouse organs, *iSEEEK* was able to identify 55 distinct cell types that are well matched with the identity and lineage of organs (**Figure 2C and Supplementary Figure 1**). In qualitative measurement of cell clustering obtained from *iSEEEK* against putative cell labels, we found that *iSEEEK* achieved an adjusted rand index (ARI) of 0.61 for HCA bone marrow dataset, 0.34 for PBMC dataset, 0.72 for Tabula Mursi dataset. The ARI metric achieved by *iSEEEK* was comparable to those achieved by Scanpy. The ARI metric and UMAP plots of Scanpy across these three datasets were provided in **Supplementary Figure 2-4**.

**Figure 2.**
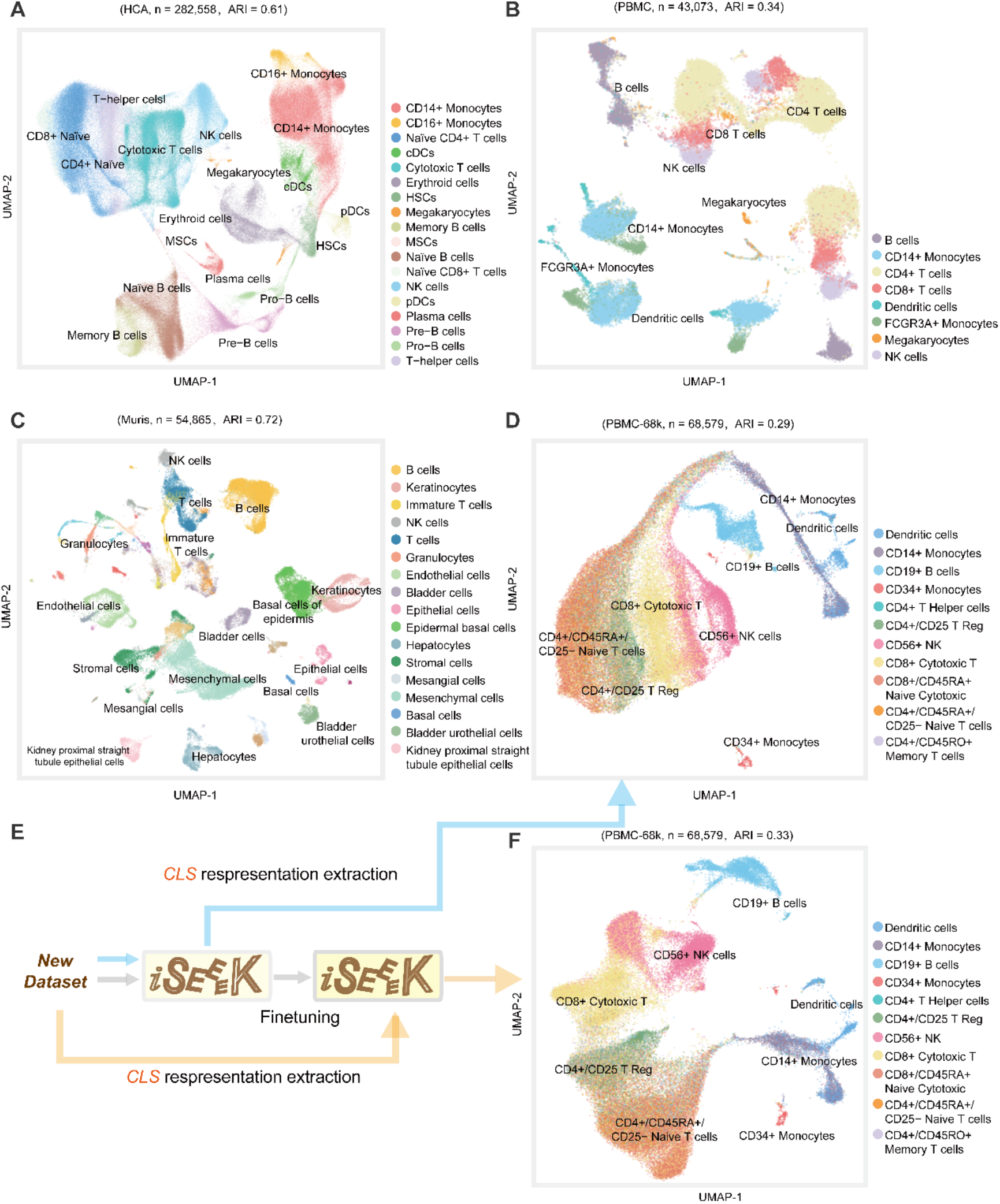
The clustering performances of *iSEEEK*. UMAP visualization of feature representations learned by *iSEEEK* on the (**A**) HCA dataset, (**B**) PBMC dataset, (**C**) Tabula Mursi dataset and (**D**) PBMC-68k dataset that was not involved in the development of *iSEEEK*. (**E**) Fine-tuning *iSEEEK* with new dataset PBMC-68k. (**F**) UMAP visualization of feature representations of PBMC-68k dataset with features extracted from *iSEEEK* being fine-tuned on the PBMC-68k dataset.

Additionally, we found that *iSEEEK* can work effectively on new dataset that was not involved in the development of *iSEEEK*. As an example, we examined *iSEEEK* on a new dataset obtained from previous study that consisted of 68,579 peripheral blood mononuclear cells from a healthy donor^23^. *iSEEEK* achieved an ARI of 0.29, which was comparable to Scanpy (**Supplementary Figure 5**), and the UMAP-visualization of the new dataset was shown in **Figure 3D**. Subsequently, we finetuned *iSEEEK* model on this new dataset (**Figure 3E**). We observed that the finetuned *iSEEEK* model achieved an ARI of 0.33 (**Figure 3F**). We found that finetuning *iSEEEK* for one epoch is sufficient (**Supplementary Figure 6**). The UMAP visualization plots of finetuning *iSEEEK* with different epochs were provided in **Supplementary Figure 6**. In addition, we showed that *iSEEEK* achieved a comparable acceptance rate of kBET as compared with batch-correction methods such as ComBat^24^, MNN^25^ and BBKNN^26^ measured on the HCA bone marrow dataset (**Supplementary Figure 7**).

**Figure 3.**
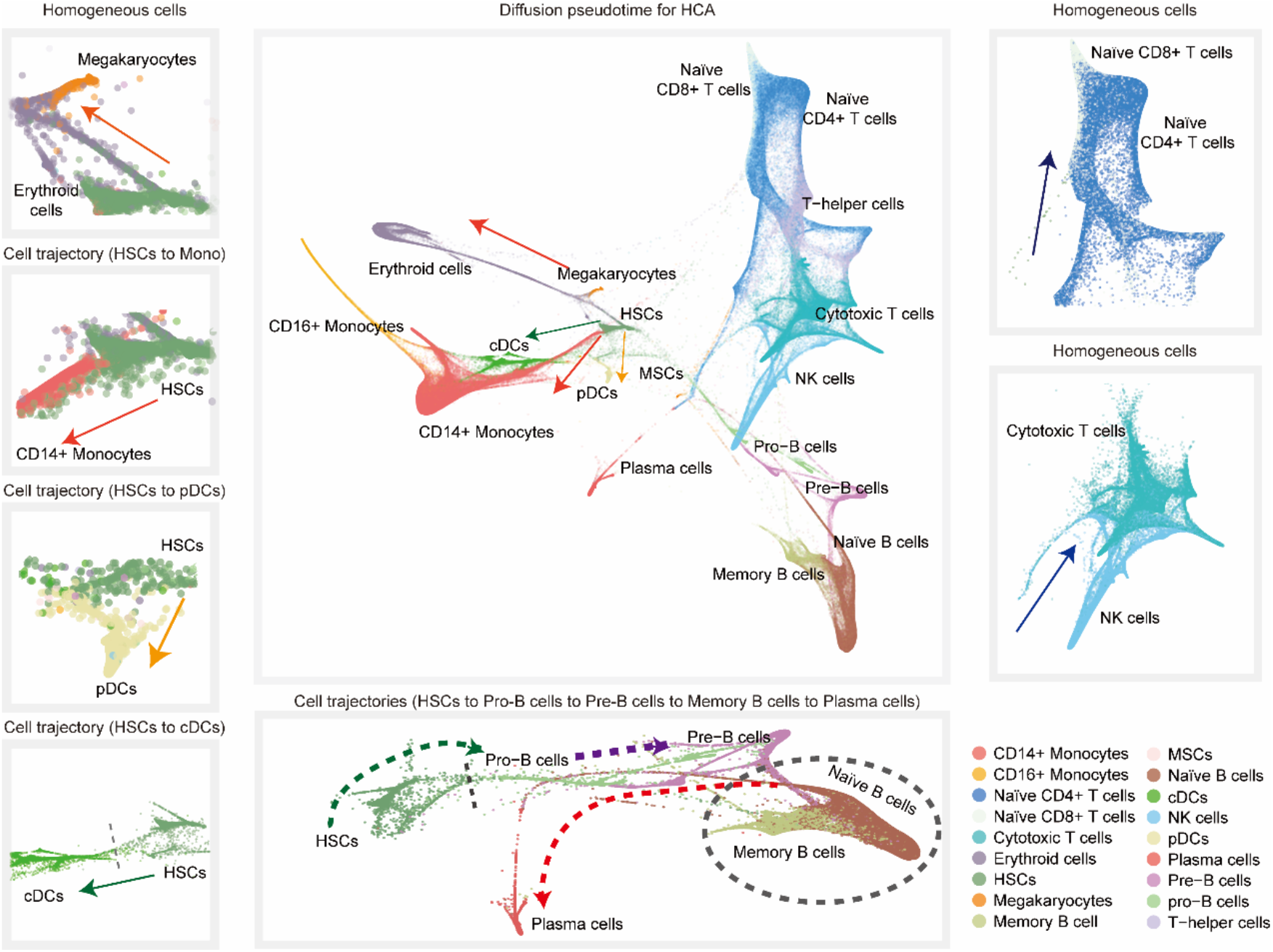
Diffusion pseudotime analysis of bone marrow cells in HCA dataset. (**A**) The panorama diffusion map of HCA dataset with the cell types colored. (**B**) Bifurcation of megakaryocytes and erythroid cells. Bifurcation of CD14+ monocytes (**C**), plasmacytoid dendritic cells (pDCs) (**D**) and conventional dendritic cells (cDCs) (**E**) from hematopoietic stem cells (HSCs). (**F**) The developmental trajectory of B cells from hematopoietic stem cells (HSCs), towards B cell progenitors (Pro-B cells), precursors of B cells (pre-B cells), matured naïve B cells, memory B cells and plasma cells. The arrows represent the directionality of the cell developmental trajectory. (**G**) Bifurcation of naïve CD4+ T cells and naïve CD8+ T cells, similarly, (**H**) cytotoxic T cells and NK cells.

### *iSEEEK* reserves the development trajectory of B cells on HCA dataset

We used the feature representation learned by *iSEEEK* to construct pseudo-temporal trajectories of bone marrow cells on HCA bone marrow dataset (see **Methods**). We identified a developmental trajectory rooted at stem cells towards multiple cell types with distinguishable intermediate stages (**Figure 3A**). We identified a developmental trajectory of B cells (**Figure 3**), with an initial wave of B cell progenitors (Pro-B cells) derived from hematopoietic stem cells (HSCs), then followed by precursors of B cells (pre-B cells), matured naïve B cells (**Figure 3F**), and finally bifurcated into memory B cells and plasma cells^27^. Meanwhile, we also observed differentiation of HSCs into multiple types of immune cells including plasmacytoid dendritic cells (pDCs), conventional dendritic cells (cDCs) and CD14+ monocytes (**Figure 3C-E**). In addition, the baicalia type of cell trajectories were observed for megakaryocytes and erythroid cells^28^ (**Figure 3B**), naïve CD4+ T cells and naïve CD8+ T cells (**Figure G**), cytotoxic T cells and NK cells (**Figure 3H**), suggesting that they were originated from the same progenitor cells^29^.

### *iSEEEK* enables discovery of marker genes and gene interaction modules

We added and trained a classifier at the end of *iSEEEK* for identification of marker genes on the dataset of FACS-sorted CD4/8+ T cells (see **Methods**). An apparent separation of CD4+ and CD8+ T cells were observed on the UMAP visualization plot (**Figure 4A**). We identified cell-type specific markers for these CD4/8+ T cells (see **Methods**). The identified marker genes for CD4+ T cells include *CD4, TXNIP* and *CD2* (**Figure 4B**). CD8+ T cells were featured by cytotoxic markers such as *CD8A, CD8B, KLRK1* and *NKG7*(**Figure 4B**).

**Figure 4.**
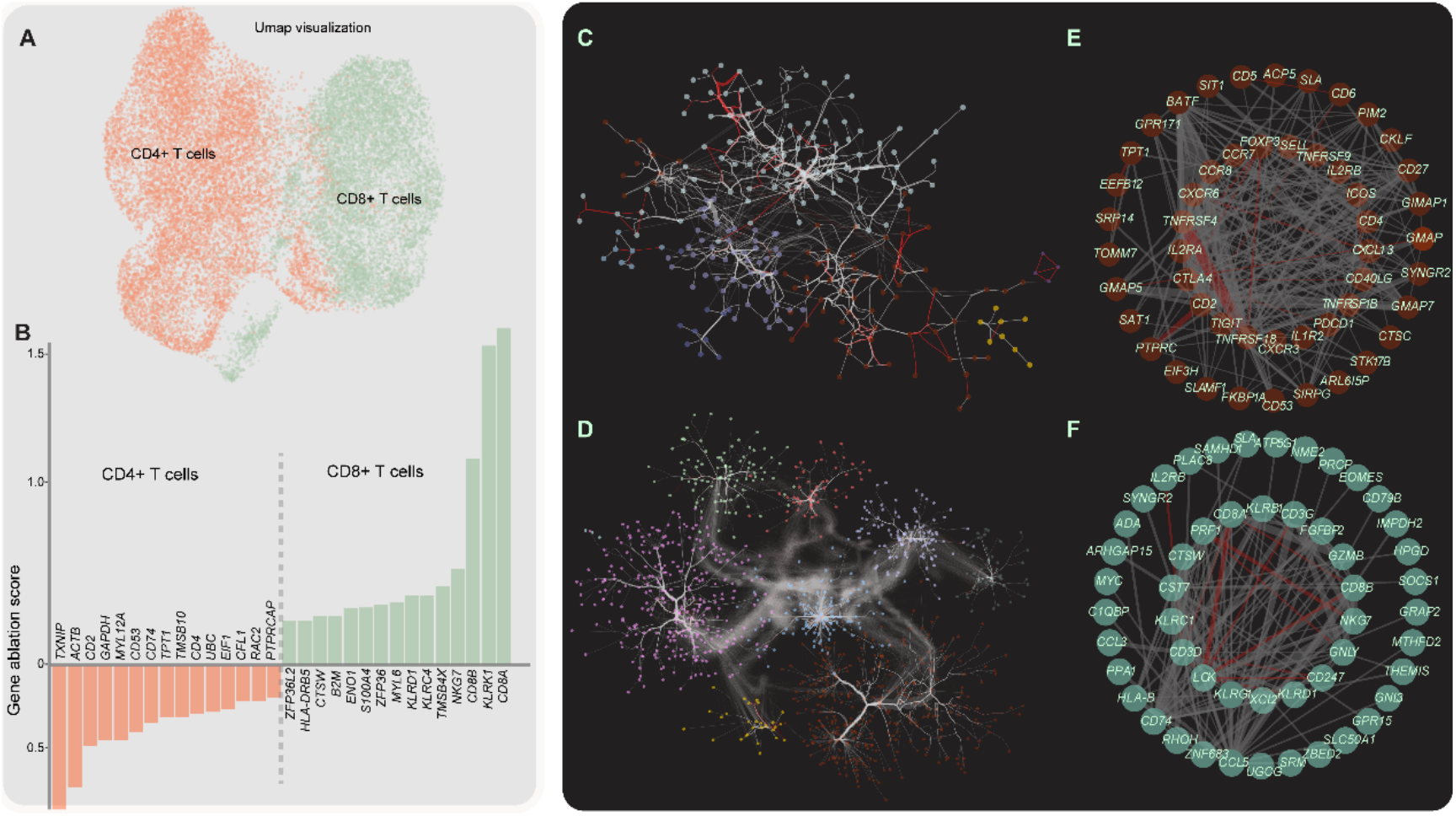
Marker genes and examplified gene-gene interaction networks deciphered from FACS-sorted CD4/8+ T cells dataset. (**A**) UMAP visualization CD4+ and CD8+ T cells. (**B**) Barplot representation of marker genes for CD4+ and CD8+ T cells. (**C and D**) The gene-gene interaction networks for CD4+ and CD8+ T cells, respectively. (**E and D**) The gene interaction modules characteristic of CD4+ and CD8+ T cells, respectively. The red edge indicates it is represented in STRING gene-gene interaction database. The thickness of the edge is proportional to attention weights among interacted genes.

We respectively obtained gene interaction networks that are characteristic of CD4+ and CD8+ T cells through analyzing the attention matrices of *iSEEEK* for the dataset of FACS-sorted CD4/8+ T cells (See Methods, **Figure 4C and 4D**). A CD4+ T cell specific gene interaction module (**Figure 4E**) derived from **Figure 4C** was featured by genes that involved in the development and function of CD4+ T cells (i.e. *CXCR6, FOXP3, ICOS, CCR7* and *SELL*)^30^ and immune suppression (i.e. *PDCD1, TIGIT, BATF* and *TNF* receptor family) ^31-33^ (**Figure 4E**). These interactions are overrepresented in the STRING gene-gene interaction database (16/244 interactions; hypergeometric test, *p* = 5.0e-4). Among these interactions, *CD2/PTPRC* interaction is involved in the activation of T cell receptor^34^. *FOXP3/TNFRSF18* interaction is critical for T cell differentiation^35^. The CD8+ T cell specific module (**Figure 4F**) is characterized by interactions among cytotoxic genes including *GNLY, NKG7, PRF1, LCK* and *KLRD1*^36^. In addition, the CD8+ T cell recruitment gene *CCL5*^37^ exhibited strong interaction with markers of CD8+ T cells including *CD8A, CD8B* and *GZMB*. Gene interactions from the CD8+ T cell specific module is enriched in STRING database (12/144 interactions; hypergeometric test, *p* = 1.3e-3).

## Discussion

In this study, we presented a universal approach *iSEEEK* for integrating super large-scale single-cell transcriptomes by exploring of the rankings of top-expressing genes. *iSEEEK* was developed on 13,702,899 single-cell transcriptomes covering a wide variety of cell-types from *Homo sapiens* and *Mus musculus*. The notable features of *iSEEEK* is that it only relies on gene rankings but not actual expression levels, thus its sensitivity to batch effect should be decreasing. This feature makes *iSEEEK* a good candidate for integrating super large-scale amount of single-cell expression data. The performance of *iSEEEK* is expected to improve as more and more data are involved in its development.

This study demonstrated that pretraining on the rankings of top-expressing genes from super large-scale scRNA-seq data is effective. The efficiency of cell cluster delineation on the extracted latent features of the pretrained *iSEEEK* was demonstrated on three heterogeneous datasets encompassing different cell types, sequencing with different protocol and deriving from different species. Across these three datasets, *iSEEEK* achieved comparable ARI metric as compared with Scanpy. In addition, *iSEEEK* also worked efficiently on new dataset that was not involved in its development. Finetuning *iSEEEK* for one epoch apprears sufficient to improve its clustering performance.

*iSEEEK* enables to maximize the value of big data from single-cell transcriptomes in simple and yet effective way. *iSEEEK* can make use of single-cell transcriptomes from different species, which was exemplified by the integration of data from *Homo sapiens* and *Mus musculus* in our study. *iSEEEK* circumvents the tremendous challenge of batch-correction in single-cell integration by modeling gene expression rankings rather than actual expression levels. As *iSEEEK* is not relying on actual expression levels but rather on the ranking of top-expressing genes, its sensitivity to batch effect is decreasing, which was verified in this study (**Supplementary Figure 7**). Batch-correction methods such as ComBat^24^, MNN^25^ and BBKNN^26^ require explicit knowledge of the batch information. However, the batch information is not always available and often neglected by researchers; therefore, traditional methods are not appropriate for data integration of multiple datasets without batch information. In addition, traditional methods^8,9^ are memory hungry as they require to load all data into memory, hampering their ability to process super large-scale dataset. In contrast, *iSEEEK* was trained in a stochastic manner that only a small batch of samples are processed at each time step. Thus, memory consumption of *iSEEEK* is much lower than traditional methods and it can benefit from acceleration brought by graphical processing unit.

*iSEEEK* is quite different from that of other traditional methods as they require selection of hyper-variable genes (HVGs), batch-correction and data normalization^38,39^, whereas *iSEEEK* uses the ranking of top-expressing genes and does not require selection of HVGs. Batch-correction methods are sensitive to data volume and the number of batches, and the robustness of the batch-correction is difficult to assess in large-scale dataset^24-26^. Meanwhile, the consistency and reproducibility of the HVGs is also difficult to control by different HVG selection methods^13^. *iSEEEK* takes as input the rankings of top-expressing genes, which may be less informative intuitively as compared with the use of expression levels of HVGs as traditional methods. However, *iSEEEK* was able to precisely identify cell types of small proportions such as FCGR3A+ and CD14+ monocytes in the PBMC dataset (**Figure 2B**), suggesting that the rankings of top-expressing genes are sufficient for delineation of cell types with small proportions.

We demonstrated that feature representation of the rankings of top-expressing genes learned by *iSEEEK* preserved the chronological order of cell development trajectories. We verified the continuous and identifiable cell trajectory from B cell progenitors derived from HSCs towards plasma cells^27^ on HCA bone marrow dataset (**Figure 3F**).

As a preliminary endeavor, we demonstrated that by analyzing *iSEEEK* for the input of CD4/8+ T cells, we were able to identify gene interaction modules manifested the features of CD4/8+ T cells. Functional related tend to have strong interactions. The attention mechanism in *iSEEEK* makes it possible to learn interaction among different genes. As the attention mechanism enables modeling gene interaction by taking into account the influence of other genes, it has the potential to learn complex gene-gene interaction networks and may shed new lights on gene regulation circuits.

In this study, we formulate single-cell transcriptome integration as a language modeling task. Recent advances in natural language processing will benefit single-cell integration. The paradigm of pretraining-then-finetuning is a de facto procedure in natural language processing as this paradigm is robust to overfitting and has the advantage of making use of super large-scale data and reducing the need of big data on downstream tasks^40^.

Herein, we provided a universal, scalable, transferable, effective and easy-to-use approach for integration of super large-scale single-cell transcriptomes. *iSEEEK* can be finetuned on a specific dataset to tackle specific downstream tasks. We expected that *iSEEEK* may be helpful for researchers to elucidate the heterogeneous and dynamic biological processes underlying human diseases with the accumulation of single-cell transcriptomes.

## Conclusions

In the study, we presented a universal approach for integrating super large-scale for single-cell transcriptomes by modeling feature representation of the rankings of top-expressing genes as a masked language modeling task. We are in the process of developing a web server running *iSEEEK* that would be freely available to the research community. Our work represented a new paradigm in the integration of super large-scale single-cell transcriptomes and may be helpful for the elucidation of the dynamic and heterogeneity of single-cells.

## ACKNOWLEDGEMENTS

We are grateful for researchers for their generosity to made their data publicly available. This work was supported by the National Natural Science Foundation of China (no. 31801117 to X.L., no. 31900471 to M.Y. and 82073287 to Q.Z.), Tianjin Municipal Health Commission Foundation (grant no. RC20027 to Y.L) and IRT_14R40 from the Program for Changjiang Scholars and Innovative Research Team in University in China (K.C.).

## AUTHOR CONTRIBUTIONS

Xiangchun Li and Kexin Chen designed and supervised the study; Hongru Shen, Xilin Shen, Mengyao Feng and Xiangchun Li performed data collection, analysis, and wrote the manuscript; Hongru Shen, Xilin Shen and Xiangchun Li developed the model; Chao Zhang, Dan Wu, Xilin Shen, Mengyao Feng, Jiani Hu, Jilei Liu, Yichen Yang, Yang Li, Meng Yang, Wei Wang and Qiang Zhang collected data; Xiangchun Li, Kexin Chen, Jilong Yang and Hongru Shen revised the manuscript.

## DECLARATION OF INTESTS

The authors declare that they have no conflict of interest.

## Methods

### Dataset and preprocessing

We collected expression matrices of 13,702,899 single-cells from previous studies. Detailed information for these studies were provided in **Supplementary Table 1**. We discarded mitochondrial genes, ribosomal genes and non-protein coding genes. Subsequently, we concatenated the 126 top-expressing genes with CLS and SEP tokens as a sentence for each single-cell. Eventually, we obtained a text file of 13,702,899 sentences. The five datasets used in downstream task of *iSEEEK* were described below:

#### Human Cell Atlas

Bone marrow data of 282,588 cells from 64 healthy donors in HCA project subjected to 10x sequencing protocol^21^. There are 18 cells types annotated by HCA including erythrocytes, mesenchymal stem cells, hematopoietic stem cell and diverse immune cells.

#### Peripheral Blood Mononuclear Cells (PBMC)

This dataset was download from Gene Expression Omnibus repository^9^ (GSE96583). It consists of 43,095 single cells obtained from 5 individuals (3 systemic lupus erythematosus and 2 control) subjected to 10x sequencing. All cells were grouped into 8 categories: B cells, CD4+ T cells, CD8+ T cells, dendritic cells, megakaryocytes, FCGR3A+ monocytes, CD14+ monocytes and natural killer cells.

#### Tabula Mursi

A data set of 100,000 single-cell from Mouse Cell Atlas^5^ across 20 different organs subjected to 10x and Smart-seq2 sequencing protocols. 54,865 cells were sorted by FACS, therefore, we used these 54,865 cells for evaluation.

#### Peripheral Blood Mononuclear Cells-68k (PBMC-68k)

This PBMC-68k dataset included 68,579 peripheral blood mononuclear cells obtained from a healthy donor (http://support.10xgenomics.com/single-cell/datasets).

#### FACS-sorted CD4/8+ T cells

This dataset includes 12,670 CD4+ and 9,012 CD8+ T cells that were sorted by FACS from tumor patients diagnosed with liver cancer, colorectal cancer and lung cancer^41-43^. They were subjected to smart-seq sequencing.

### The *iSEEEK* model

*iSEEEK* consists of an embedding layer and 8 encoder layers each with 576 hidden units and 8 attention heads.

#### Embedding Layer

The embedding layer takes the embeddings of a sequence of 128 tokens and their position embeddings as input. An input representation of token can be represented as [*CLS,G*_1_,*G*_2_, …,*G*_*n*_, *SEP*]. *CLS* is the classification token and *SEP* is sentence separation token. *G*_*i*_ is the gene symbol of the i^th^ gene. The CLS token, gene symbols and SEP token are first converted into indexes in the gene symbol dictionary. The gene symbol dictionary consists of protein-encoding genes.

#### Encoder layer

The encoder layer is a transformer that is the core component of *iSEEEK*. It consists of a multi-head self-attention and a feed-forward network inter-connected with layer normalization layer. Residual connection is added to improve information flow. The multi-head self-attention enables the model to capture contextual information. The self-attention head is formulated as:

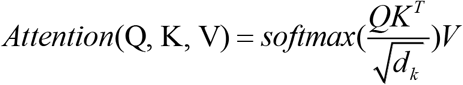

The self-attention head takes *Q, K* and *V* as inputs and applies softmax transformation. *Q, K* and *V* are projected from the input. The scaling factor 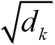 is used to mitigate the extreme small gradient^44^.

### Input representations

We constructed a dictionary with protein-encoding genes. For each cell, we prepared a sequence of 128 tokens, where tokens are gene symbols and/or special tokens such as [CLS], [SEP] and [PAD]. We filtered out genes with extremely low expression (i.e. an expression level of 1 or 0) and ranked them according to their expression levels. We padded [PAD] token to the input sequence if the number of genes is less than 126. The first token is always [CLS] and the last token is always [SEP].

### Model pre-training

*iSEEEK* take a sequence gene symbols with a maximum length of 126 as input. We applied the same data sampling strategy during training as BERT^19^: the training data generator randomly chooses 15% of the gene positions for prediction. If the *i*^*th*^ gene is chosen, we replace the it with (1) the [MASK] token 80% of the time, (2) a random gene 10% of the time, (III) the original unchanged gene 10% of the time. *iSEEEK* was trained by cross-entropy loss by comparing its predictions to the original genes. We trained *iSEEEK* model for 48 epochs with a batch size of 64 and the learning rate was set to 0.0001. The *PyTorch* (version 1.7.1) and *transformers* (version 4.6.0) packages were used to develop *iSEEEK*.

### Identification of marker genes

We added a classifier to the end of the pre-trained *iSEEEK* and trained on the FACS-sorted CD4/8+ T cells. The parameters of the pre-trained *iSEEEK* were frozen and parameter updating was applied for the linear classifier. We trained this classifier with a learning rate of 0.001 and batch size of 16 with Adam optimizer for 30 epochs. We quantitatively measure the impact of a specific gene as the difference between the logit values for the original gene sequence and gene sequence with that gene replaced with [UNK] token. Specifically, for an input gene sequence of ***S*** = [*G*_*1*_, *G*_*2*_, …, *G*_*n*_], we obtained ***S***^***\****^ = [*G*_*1*_, *UNK*, …, *G*_*n*_] by replacing *G*_*2*_ with *UNK*. Let *L* and *L*^*^ denote the logit values obtained from the classifier, the influence of *G*_*2*_ on the decision made by this classifier is defined as:

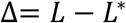

For a specific cell type, we rank the influence of genes by the average value of Δ and those ranked on the top is considered to be marker genes.

### Diffusion pseudotime analysis

The affinity matrix of cells *W*_*n*×*n*_ was constructed from representation features of the *CLS* token. which is performed using community detection algorithms^45^ and the HNSW algorithm^46^ is applied to find the top-k nearest neighbors. A scaled Gaussian kernel is used to define the distance between cell-*x* and cell-*y* as:

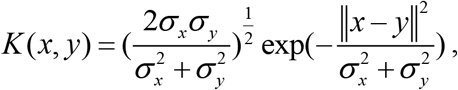

*x* and *y* are representation features of the *CLS* token for cell-*x* and cell-*y*, respectively. *σ* _*x*_ is the local kernel width of *x*, calculated as the median value of *x* and its top-k nearest cells. The affinity matrix is defined as:

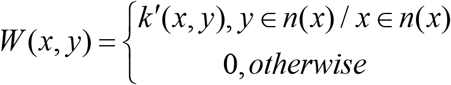

Where *k*′(*x, y*) is defined as:

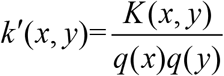

The Markov chain transition matrix *P* and the symmetric transition matrix *Q* are then calculated based on the affinity matrix as follows:

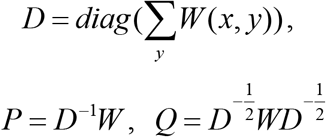

The symmetrical matrix *Q* can be decomposed as *UAU*^*T*^. Let 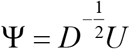. A family with parameter timescale of *t* for approximated diffusion maps {Ψ_*t*_}_*t*∈•⋃{∞}_ is defined as:

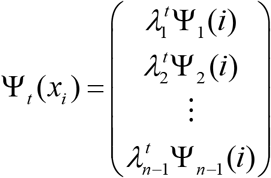

The approximated DPT maps 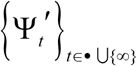 are constructed based on the aforementioned diffusion maps as:

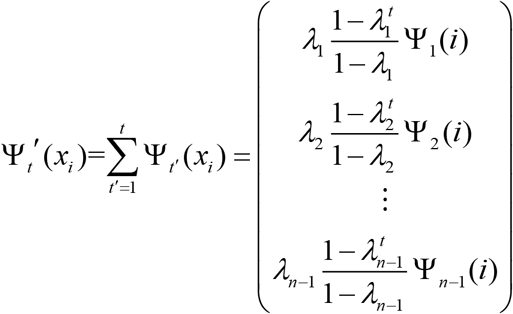

The diffusion maps and diffusion pseudotime maps are performed using package *Pegasus*^47^ (v1.4.3) with K set to 30. The cell trajectory was visualized with force-directed layout embedding (FLE) algorithm^48^. We set δ and nδ as its the default parameter: δ=2.0 and nδ=5,000.

### Construction of gene interaction network

We constructed the cell-type specific gene interactions respectively for CD4+ and CD8+ T cells based on the FACS-sorted CD4/8+ T cell dataset^23^. For each input sequence consisted of *n* genes, we can extract an attention matrix ***a*** of *n* columns and *n* rows corresponding to each attention head. Attention weight *a*_*i,j*_ denotes the attention of gene *i* to gene *j*. Gene attention matrix of a specific cell type was constructed from the attention matrix ***a*** for each cell from that cell type. Specifically, we define an indicator function *f*(*i, j, θ*) that returns 1 if the attention weight between gene *i* and *j a*_*i,j*_ > *θ*, and 0 otherwise. The attention matrix a specific cell type (*C*_*a*_) was constructed as follow:

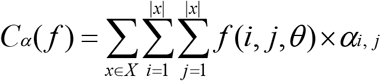

*θ* is a threshold to filter out low attentions and a value of 0.05 was used in this study. Given that attentions between gene *i* and *j* is not identical to *j* and *i*, therefore, the attention matrix a specific cell type was further refined as:

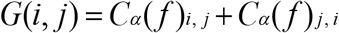

We retained the top 10% interactions in *G(i, j)* in subsequent analysis. Network construction was carried out with Python package *networkx* (version 2.5). Functional modules of networks were detected through Louvain community detection algorithm^49^ based on package *python-community* (version 0.15). Overrepresentation of detected modules in STRING gene-gene interaction database^50^ was evaluated with hypergeometric test. A *p* < 0.05 was considered statistically significant. The gene interaction networks were visualized using *Cytoscape* (version 3.8.2)^51^.

### Single-cell clustering and evaluation

We extracted the represented features of each single-cell with the pretrained *iSEEEK*. The extracted features were used as input to the K-Nearest Neighbors (KNN) algorithm to construct KNN graphs for subsequent single-cell community detection by Leiden^52^ algorithm. We applied single-cell clustering pipeline implemented in Scanpy to perform single-cell clustering on KNN graph. The uniform manifold approximation and projection^53^ (UMAP) is used for visualizing clustering result.

For comparison, we also performed single-cell clustering using Scanpy (v1.6.0) as the benchmarking tools. The conventional single-cell analysis based on the gene expression. We first filtered out cells and the criteria: the number of expression genes <200 or mitochondrial counts >30%. The highly variable genes (HVGs) were selected with default parameters (i.e *max_mean*=3 and *min_mean*=0.0125). We used the default 50 principal components to construct the KNN graph and subsequently applied Leiden community detection algorithm to delineate cluster with default parameter (i.e. resolution =1).

We used adjusted rand index (*ARI*) as clustering measure to evaluate the clustering performance. The *ARI* metric is calculated on the contingency table summarizing the truth labels and clustering. In the contingency table, rows and columns represent truth and clustering labels, respectively. *ARI* is defined as:

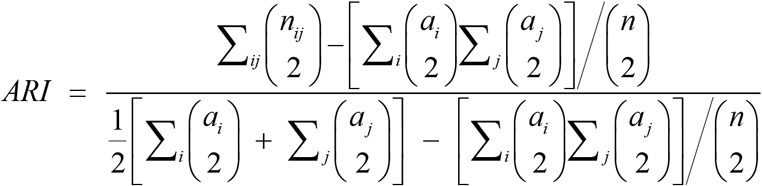

Where *n*_*ij*_ denoted the numbers of cell in common between clustering labels and truth labels, *a*_*i*_ the sum of *i*^*th*^ row and *a* _*j*_ the sum of *j*^*th*^ column of the contingency table.

### Batch-correction and evaluation

We used the acceptance rate of kBET^54^ as a measurement of batch-effect. The acceptance rate measures whether cells from different batches are well-mixed in the local neighborhood of each cell. The acceptance rate obtained from *iSEEEK* was compared with the other three batch-correction methods Combat^55^, MNN^25^, BBKNN^26^. ***kBET acceptance rate***. We assumed that the dataset of single-cell with batches of *m*, and there are *n*_*j*_ cells in batch *j*. The batch mixing frequency denotes as *f* = (*f*_1_,…, *f*_*m*_), where 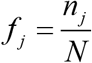. The number of neighbors of cell-*i* belonging to batch *j* is 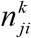. Its *χ*^2^ test statistic with degrees of (m-1) is calculated as: 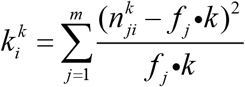. The *P* value is calculated as: 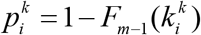, where *F*_*m*−1_(*x*) represents the cumulated density function. The kBET acceptance rate is defined as the percentage of cells that accept the null hypothesis at significance level α as follows:

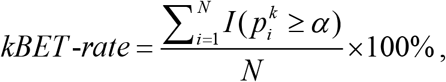

*I(x)* is the indicator function where *I(x)* = 1 if x > 0 otherwise *I(x)* = 0. We used Pegasus (v1.4.3) to calculate the kBET acceptance rate by setting *K* and α to 5 and 0.01, respectively.

## Supplementary Figures & Tables

**Supplementary Figure 1. The full annotation of UMAP visualization of *iSEEEK* on the Tabula Muris**. The ARI metric and annotation of cells are shown.

**Supplementary Figure 2. The UMAP visualization plots of Scanpy with different batch-correction methods on the HCA dataset**. Batch-correction methods included (**A**) Combat, (**B**) MNN and (**C**) BBKNN, respectively. The ARI metric and annotation of cells are shown.

**Supplementary Figure 3. The UMAP visualization plots of Scanpy with different batch-correction methods on the PBMC dataset**. Batch-correction methods included (**A**) Combat, (**B**) MNN and (**C**) BBKNN, respectively. The ARI metric and annotation of cells are shown.

**Supplementary Figure 4. The UMAP visualization plot of Scanpy on the Tabula Muris dataset**. The ARI metric and annotation of cells are shown.

**Supplementary Figure 5. The UMAP visualization plot of Scanpy on the PBMC-68k dataset**. The ARI metric and annotation of cells are shown.

**Supplementary Figure 6. The UMAP visualization plots of *iSEEEK* finetuned on the PBMC-68k dataset for 1 (A), 2 (B), 3 (C) and 4 (D) epochs, respectively**. The ARI metric and annotation of cells are shown.

**Supplementary Figure 7. The kBET acceptance rate of *iSEEEK* and Scanpy with different batch-correction methods such as ComBat, MNN and BBKNN on the HCA bone marrow dataset**.

**Supplementary Table 1. Data source information**.

